# Population dynamics hide phenotypic changes driven by subtle chemical exposures: implications for risk assessments

**DOI:** 10.1101/2023.02.02.526756

**Authors:** Ana del Arco, Lutz Becks, Inmaculada de Vicente

## Abstract

Ecological risk assessment of chemicals focuses on the response of different taxa in isolation not taking ecological and evolutionary interplay in communities into account. Its consideration would, however, allow for an improved assessment by testing for implications within and across trophic levels and changes in the phenotypic and genotypic diversity within populations. We present a simple experimental system that can be used to evaluate the ecological and evolutionary responses to chemical exposure at microbial community levels. We exposed a microbial model system of the ciliate *Tetrahymena thermophila* (predator) and the bacterium *Pseudomonas fluorescens* (prey) to iron released from Magnetic Particles (MP-Fe_dis_), which are Phosphorus (P) adsorbents used in lake restoration. Our results show that while the responses of predator single population size differed across concentrations of MP-Fe_dis_ and the responses of prey from communities differed also across concentration of MP-Fe_dis_, the community responses (species ratio) were similar for the different MP-Fe_dis_ concentrations. Looking further at an evolutionary change in the bacterial preys’ defence, we found that MP-Fe_dis_ drove different patterns and dynamics of defence evolution. Overall, our study shows how similar community dynamics mask changes at evolutionary levels what would be overlooked in the design of current risk assessment protocols where evolutionary approaches are not considered.

## Introduction

Populations are continuously exposed to biotic and abiotic changes which alter population densities as well as evolutionary responses (Matthews et al. 2011). Consumer-resource interactions are iconic examples of continuous biotic changes with often rapid fluctuations in predator and prey population sizes or continuous adaptation and, other relevant example is counter adaptation of host and parasite (Murdoch et al. 2003). Many previous studies showed how abiotic stress can promote rapid evolutionary responses and how it prevents population extinction through adaptation in response to such abiotic stress (Bell and Gonzalez 2011; Lindsey et al. 2013; Ramsayer et al. 2013; Bell 2017). The responses to both abiotic and biotic changes can however, be more complex when demographic and evolutionary changes occur simultaneously and potentially drive each other as eco-evolutionary dynamics (Yoshida et al. 2003; Post and Palkovacs 2009; Palkovacs and Hendry 2010). The significance of ecology and evolution interplay to understand community changes raises the concern about its role in ecological risk assessments, as chemical exposure can impact both ecology (e.g., population extinction and bottlenecks, species sorting within communities) and evolution (e.g., drift, fixation of alleles related to adaptation to the chemical, loss of genetic diversity, epigenetic modifications), which ultimately may feed back into each other.

Current risk assessments do not evaluate the potential for eco-evolutionary changes despite of the increasing awareness of its relevance for understanding the mechanisms driving community changes (Brady et al. 2017; De Meester et al. 2019). Assessing eco-evolutionary responses to chemical exposure is challenging because populations are part of communities where multiple species interact. Generally, responses to chemicals can lead to decreasing (e.g., chemical toxicity) or increasing population densities (e.g. competition release, fertilizer effect) depending on the concentration, taxa, and species interaction (Beketov and Liess 2006; Brooks et al. 2009; Lopez Pascua et al. 2012; del Arco et al. 2015; Sentis et al. 2017), and whether or not the effect on the interacting species is symmetrical or asymmetrical. For instance, van den Brink and co-authors (2017) concluded that the interaction between biotic (competition and predation) and abiotic (insecticide exposure) stressors on aquatic invertebrates is species- and context-specific. In addition to changes in population densities, chemical exposure can lead to altered selection on traits involved in species interactions. Finally, the ecological and evolutionary responses can influence each other (Hendry et al. 2011), making predictions on how populations and communities respond to the presence of abiotic stressors difficult. Developing these predictions are, however, important when it comes to understanding how communities respond to anthrophogenic changes (Brady et al. 2017; De Meester et al. 2019; Straub et al. 2020).

To evaluate the importance of considering evolutionary changes for risk assessment in communities, we investigate here both single species population (prey or predator) and community (predator-prey) ecological responses and population (prey) evolutionary responses to the presence of a chemical. We do this for a specific chemical release (dissolved iron) from the used of novel absorbent (Magnetic Particles, MP) in restoration of eutrophicated aquatic ecosystems. Eutrophication is mainly anthropogenically driven (Smith and Schindler 2009) and is the result of excessive phosphorus (P) loads (external and internal) causing what is ultimately observed as lake water deterioration (Jeppensen et al. 1991; Søndergaard et al. 1993; Carpenter 2008). Restoration programs aim to control P loads by using a wide range of adsorbents to remove excess P (i.e. Phoslock, Zeolite or Metsorb). Recently, Magnetic Particles (MP) have been proposed as an alternative because they have a high P adsorption capacity, fast adsorption kinetics, are pH and redox condition independent (de Vicente et al. 2010; Merino-Martos et al. 2011), and they can be reused after P-desorbing (Álvarez-Manzaneda et al., personal communication). Beside the removal of P, MP can release low to undetectable amounts of dissolved iron (MP-Fe_dis_) (Funes et al. 2016). Previous assessments observing the effect of MP application for 24 hours and MP-Fe_dis_ on lake communities (Funes et al. 2016; Álvarez-Manzaneda et al. 2019) demonstrated that a single application of MP reduced P in the water column and had no impact on the plankton communities in terms of total abundance, species richness and species diversity (Álvarez-Manzaneda et al. 2019; del Arco et al. 2021). However, in a laboratory experiment with glass jars (1 L) MP exposure effects (also 24 hours contact time) on *Daphnia magna* resulted in drastic abundance decrease independently of the degree of intra-specific competition (i.e. lower to higher densities with same food availability and space) and/or community structure (i.e. presence of absence of vegetation) (del Arco et al. 2017). These ecotoxicological studies did, however, not consider evolutionary responses to the presence of MP-Fe_dis_ and it is unclear if for instance genetic sorting within a species affected the observed community dynamics.

We tested the ecological and evolutionary responses to MP-Fe_dis_ (released from an MP concentration mimicking real filed applications) in a microbial predator-prey system with the bacterium *Pseudomonas fluorescens* as the prey and the ciliate *Tetrahymena thermophila* as the predator. We focused on microbial communities because they form the bottom of any food web and changes in abundances and interactions at this level can have significant effects on whole systems and their function (Madan et al. 2005; Worden and Not 2008). Previous work with bacteria and protists showed the effects of Fe_dis_ where population sizes decreased (Dayeh et al. 2005; Wang et al. 2014). Therefore, Fe_dis_ released from the use of MP in field applications for eutrophic restoration might impose further pressures on the community. In addition, microbial communities have short generations times, often large population sizes and thus a higher potential to evolve a response to the exposure to stressors (Bell 2017), which might increase the relative role of evolutionary change in mitigating ecological responses to chemical exposure. Simple experimental assays where one can compare evolutionary changes as changes in fitness of individuals and/or populations relative to the ancestor (Kassen 2014) will facilitate the possibility to integrate evolutionary response into risk assessment of functional changes of microbial communities.

In the *Pseudomonas* - *Tetrahymena* system, predation by the ciliates often leads to the evolution of a defence in the prey population against consumption by the predator such as biofilm formation or growth in aggregates which affects the predator-prey interaction and the relative roles of ecological and evolutionary change depending on the environmental conditions (Matz and Kjelleberg 2005; Friman et al. 2014; Hiltunen et al. 2015). In this study, we mimic an environmental MP application in the field (Funes et al. 2016) by exposing bacteria and ciliates to the MP-Fe_dis_ released from MP, we have focused on dissolved iron because MP would be retrieved from the lake after 24 hours. Specifically, treatments covered a range of MP concentrations mimicking different MP field applications: 0 g/L (control), 1 g/L and 2 g/L (hereinafter 0 MP, 1 MP and 2 MP).

## Material and methods

### Model system

We used the ciliate *Tetrahymena thermophila* 1630/1U (CCAP) as the predator and the bacterium *Pseudomonas fluorescens* strain SBW25 as the prey (hereinafter predator and prey). Predator stocks were cultured axenically in organic medium (20 g of proteose peptone and 2.5 g of yeast extract in 1 L of deionized water) prior to the experiment (Hiltunen et al. 2015). Prey was inoculated from a single colony from a frozen stock to minimize initial genetic variability in the population. Thus, evolutionary changes were expected to result from *de-novo* evolution as previously shown in this system (Hiltunen et al. 2015).

### Magnetic particles

The MP (97.5% iron, 0.9% carbon, 0.5% oxygen, and 0.9% nitrogen) were kindly supplied by BASF (Germany). MP are spherical and polydisperse with an average size range of 805 nm ± 10 nm and a density of 7.5 g cm^-3^ (de Vicente et al. 2010). Maximum P adsorption capacity has been reported as 1 g MP:18.8 mg P (de Vicente et al. 2010).

### Microcosm experiment: single populations and community dynamics

Ecological and evolutionary dynamics of predator-prey single populations and communities were followed over time in semi-continuous cultures. Our experimental design aimed to assess the interplay between ecology and evolution in the short-term (e.g. *P. fluorencens* could rapidly evolve (Friman et al. 2014)), which may have the potential to lead to eco-evolutionary feedbacks in the long term at community levels (De Meester et al. 2019).

Single populations and predator-prey communities were exposed to an initial 24 hours exposure of MP concentrations (see experimental details below). We set up four replicates with only (i) predator or (ii) prey and (iii) predator and prey together. Flasks were inoculated with 100 µL from an overnight culture of bacteria stock and 3,850 predators (Hiltunen et al. 2015) either alone (single population treatments) or together (community treatments).

We used tissue culture flasks (40 mL) supplied with 11 mL of M9 salts and King’s B (KB) nutrients [0.05x; 200 mL of Minimal Salts (5x), 20 g of proteose peptone and 10 g of glycerol for 1 L of medium]. We used three different stocks of the medium with different concentrations of MP; 0 g/L (control), 1 g/L and 2 g/L (hereinafter 0 MP, 1 MP and 2 MP) which simulated a realistic environmental MP application (Funes et al. 2016). For this, we prepared a stock solution of MP by diluting 1 or 2 g of MP in 20 mL of 0.05x KB medium. After 24 hours, MP were removed from the stock solution using a magnetic gradient (Block magnet 40×40×20 mm, Webcraft GMbH, Germany) and the medium with the dissolved iron over 24 hours (MP-Fe_dis_) was autoclaved (121 °C for 20 minutes) to establish sterile testing conditions.

For the single and the community experiment, we transferred 10% of the volume from experimental flasks every 24 hours to re-seed new flasks with fresh medium without any more MP exposure. The single population community, transfers were done during 2 (48 hours) days. And, for the community experiment for 7 days (168 hours), covering approximately 21 predator and 42 prey generations. Flasks were kept at 28º C ± 2 ºC under static conditions. At each transfer, a 2 mL subsample from each flask was frozen to cryopreserve the prey after adding 0.5 mL of 80% glycerol to the sample and stored at −80 °C until further use. Another 300 µL were taken to measure prey density using Optical Density (OD) at 600 nm (Tecan absorbance microplate reader). OD measurements were transformed to *prey* Colony Forming Units (CFU). A sample of ancestral prey and experimental samples (n=3) grown overnight, later diluted, platted on solid media and we determined CFUs to establish the conversion between OD and CFU/mL. In addition, another 1 mL subsample was fixated with formaldehyde (4% final concentration) for later estimation of predator density using flow cytometry (FACS Calibur). We used the auto sampler (3 × 10 μl samples) in the flow cytometer using well plates. Prior to flow cytometer, samples were stained with 1 μl of 100x concentration of SYBER Green I (Sigma-Aldrich), and incubated for 1 hour at 24°C in the dark (Ayan 2018). Prey and predator densities were used to calculate growth rates, predator-prey ratios, and follow population dynamics.

### Fe measurements

We prepared additional culture flasks to measure MP-Fe_dis_ from each MP treatment without the addition of prey or predators. From these suspensions, samples were taken at different times: 0 hours (right after the addition of the MP) and after 24 hours (after removing the MP). Total MP-Fe_dis_ was measured using the spectrophotometric ferrozine method (Gibbs 1979). MP-Fe_dis_ was measured within 24 hours to mimic environmental MP application in the field where MP would be removed after 24 hours. In addition, in the microcosm experiments, 90% of medium was renewed with unconditioned medium every 24 hours after the initial 24-hour exposure of the populations and community.

### Evolutionary changes in prey populations

The evolution of a prey defence against predator grazing was estimated by using a simple, ecologically relevant bioassay described in detail in (Hiltunen and Becks 2014). After thawing cryopreserved prey populations taken from the start and at every transfer and growing populations for ∼10 generations (24 hours) in liquid culture (0.05% KB), 100 μL of the culture was added to 2 mL of fresh culture medium. 3,850 predators were added from the predator stock culture (i.e., naïve predators as a standard for consumer feeding on potentially genetically differentiated prey) and predator growth rate was estimated over 48 hours. Differences in predator densities compared to predators’ growth of naïve prey were taken as an estimate of the prey defence level (D). Prey defence trait values were calculated as relative fitness by D = 1-(prey_evo_/prey_anc_) where prey_evo_ is the predator density after feeding on evolved prey, and prey_anc_ is predator density after feeding on ancestral prey (coming from frozen cultures, same strain used to start both populations and community treatments).

### Statistical analysis

All statistical analysis were done with R 4.1.1 (Development Core Team). Differences in MP-Fe_dis_ concentrations between control and treatments were assessed by using a non-parametric Wilcoxon Rank Sum test, using the PMCMR package (Pohlert 2014). We used Generalized linear model (GLM), using the geepack package (HÃjsgaard et al., 2006) for assessing differences between predator and prey densities in both prey and predator coming from single populations or communities. For single population, we compared densities between MP treatment (0 MP, 1 MP, 2 MP) using time as factor. And for populations coming from communities, we assessed density differences between MP treatment and predation treatment (prey *vs*. prey under predation). Also, we used Generalized Estimating Equation models (GEE), using the geepack package for assessing differences between defence levels, predator-prey ratios over time with MP treatments and time as fixed effects. The GEE models correct for autocorrelation between time points within one replicate and we used independence correction (Halekoh et al. 2006). For data analysis, different models were used depending on the kind of data; family gamma for predator densities, family gamma for prey densities, and family poisson for predator-prey dynamics, family gamma for ratios and family gaussian for defence levels. We used species-specific models (prey or predator) to test the effects of MP-Fe_dis_ in the single species treatments and models using only data from the predator-prey treatments to test for the effect of MP-Fe_dis_ on predator-prey ratios and defence evolution (Hiltunen and Becks 2014; Hiltunen et al. 2015).

## Results and discussion

### Effects of MP-Fe_dis_ on prey and predator growth

There were significantly different amounts of MP-Fe_dis_ released in the 2 MP treatments with respect to the 1 MP and 0 MP control (Kruskal-Wallis: chi-squared = 10; p-value < 0.01; right after removing of the MP, Kruskal-Wallis: chi-squared = 10; p-value < 0.01, Table 1). Even these low amounts of iron during MP application) have the potential to impact microbes over multiple generations (i.e. ∼8 to 16 generations within 24 hours for bacterium taxa as Escherichia, Pseudomonas, Salmonella, Staphylococcus or Vibrio (Gibson et al. 2018)). Detrimental effects of metals on microbes have been previously reported (Madoni and Romeo 2006; Wang et al. 2014). The lethal concentration (LC_50_) for Fe_dis_ is ∼ 7 mg/L - 4 mg/L (in alamar Blue and CFDA-AM media) for the predator *T. thermophila* (Dayeh et al. 2005). The minimum inhibitory concentration (MIC) for the prey *Pseudomonas* has been reported as ≤ 0.349 mg/L (Workentine et al. 2008; Chiadò et al. 2013). The MP-Fe_dis_ in our study was lower than reported LC_50_ and on the range of MIC. These low concentrations might not impose a direct risk or toxic effect to higher biological levels, but might impact the microbial loop and its role for nutrient cycling as well as higher trophic levels. For instance, changes in the bacteria and ciliates communities might influence food quality for higher trophic levels. Such changes in food quality for grazers as Cladocera do not only impact growth and grazing efficiency but can trigger changes in reproduction cycles (sexual *vs*. asexual) and resting eggs production (Koch et al. 2009).

**Table 1.**
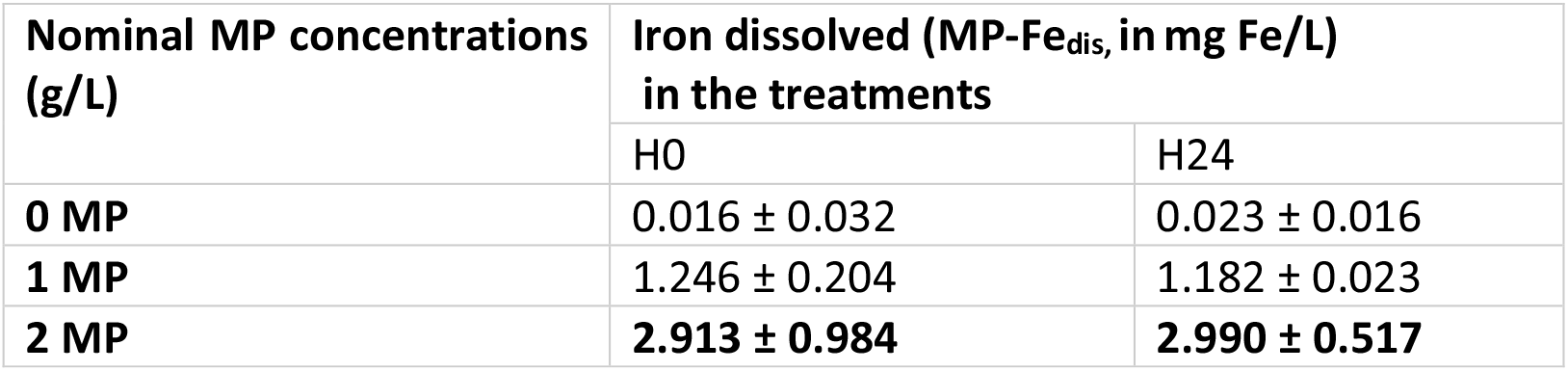
Temporal changes of dissolved iron (MP-Fe_dis_) concentrations (mean ± s.d., *n*=4) measured right after application of MP (H0) and after 24 hours (H24). Numbers in bold indicate significant differences (*p*<0.05) with respect to the iron released (Fe_dis_) from magnetic particle concentrations compared to the controls (0 MP).

We used single populations with only prey or predator to test for the effects of the MP-Fe_dis_ from the MP treatments on the two species separately for 48 hour exposure. Average predator densities significantly differed with increasing MP concentrations but were did not change between over time (Generalized linear model (GLM), MP: F=5.5133, df=2, p=0.0136; Timer: F=3.3997, df=, p=0.0817; Time*MP: F=0.304, df=1, p=0.7413; Fig. 1a).

**Fig. 1.**
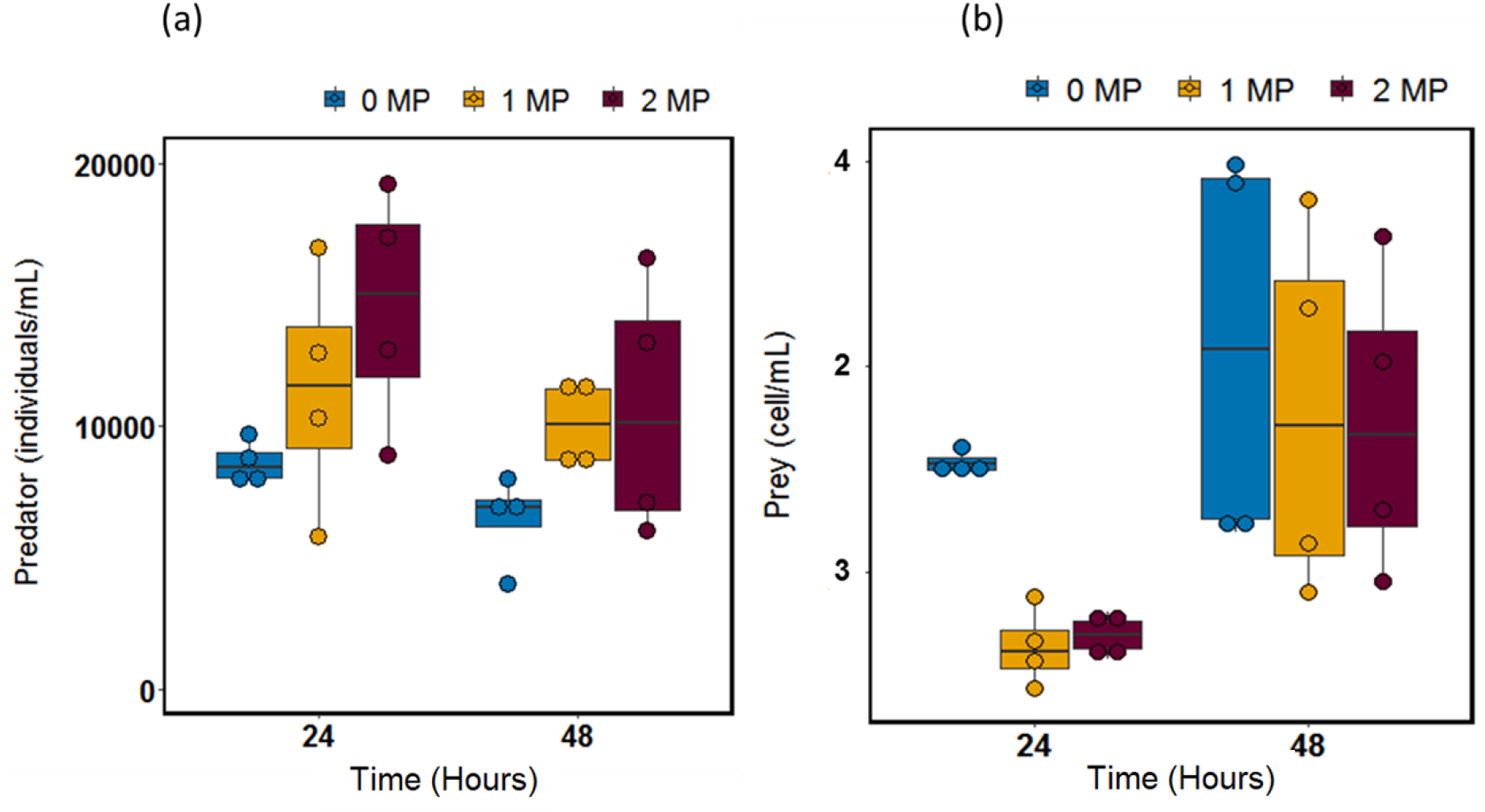
Predator (a) and prey (b) population size (10^9^) changes in single populations (predator or prey growing alone, median, n=4) in the absence and presence of MP-Fe_dis_ released from the three magnetic particles concentration tested (0 MP, 1 MP and 2 MP).

For the prey populations, we observed that the average prey densities were significantly different when comparing start and end of the experiment (GLM, Time: F=17.2743, d=1, p=0.00059) but only marginally across MP concentrations (MP: F=3.1124, df=2, p=0.069, MP*Time: F=2.2286, d=1, p=0.137; Fig. 1b).

Overall, the MP-Fe_dis_ released from MP concentrations (intended to be used in field applications) had an asymmetric effect on the species. MP-Fe_dis_ might have altered the predator populations by acting as a micronutrient right after exposure, facilitating the growth of predator (Fig. 1a). On the contrary, MP-Fe_dis_ negatively affected the bacteria resulting in smaller population size compared to the controls right after exposure (Fig. 1b), however, bacteria reached similar densities after 48 hours in all treatments what might suggest adaption to the exposure (e.g. genotype sorting, *de-novo* mutations).

### Effects of MP-Fe_dis_ on predator-prey interaction

When prey and predator grew together, predators significantly reduced prey densities depending on the MP concentration over time (GLM, MP: F=227463702, df=2, p<2×10^−16^; Predation: F=18121171, df=1, p<2×10^−16^; MP x Predation: F=0, df=2, p<2×10^−16^; Fig. 2 “All experimental days”), with increasing prey densities with increasing MP concentrations under predation conditions, specially by the end of the experiment (GLM, MP: F=1330690697, df=2, p<2×10^−16^; Predation: F=603799903, df=1, p<2×10^−16^; MP x Predation: F=388377847, df=2, p<2×10^−16^; Fig. 2, Last experimental day).

**Fig. 2.**
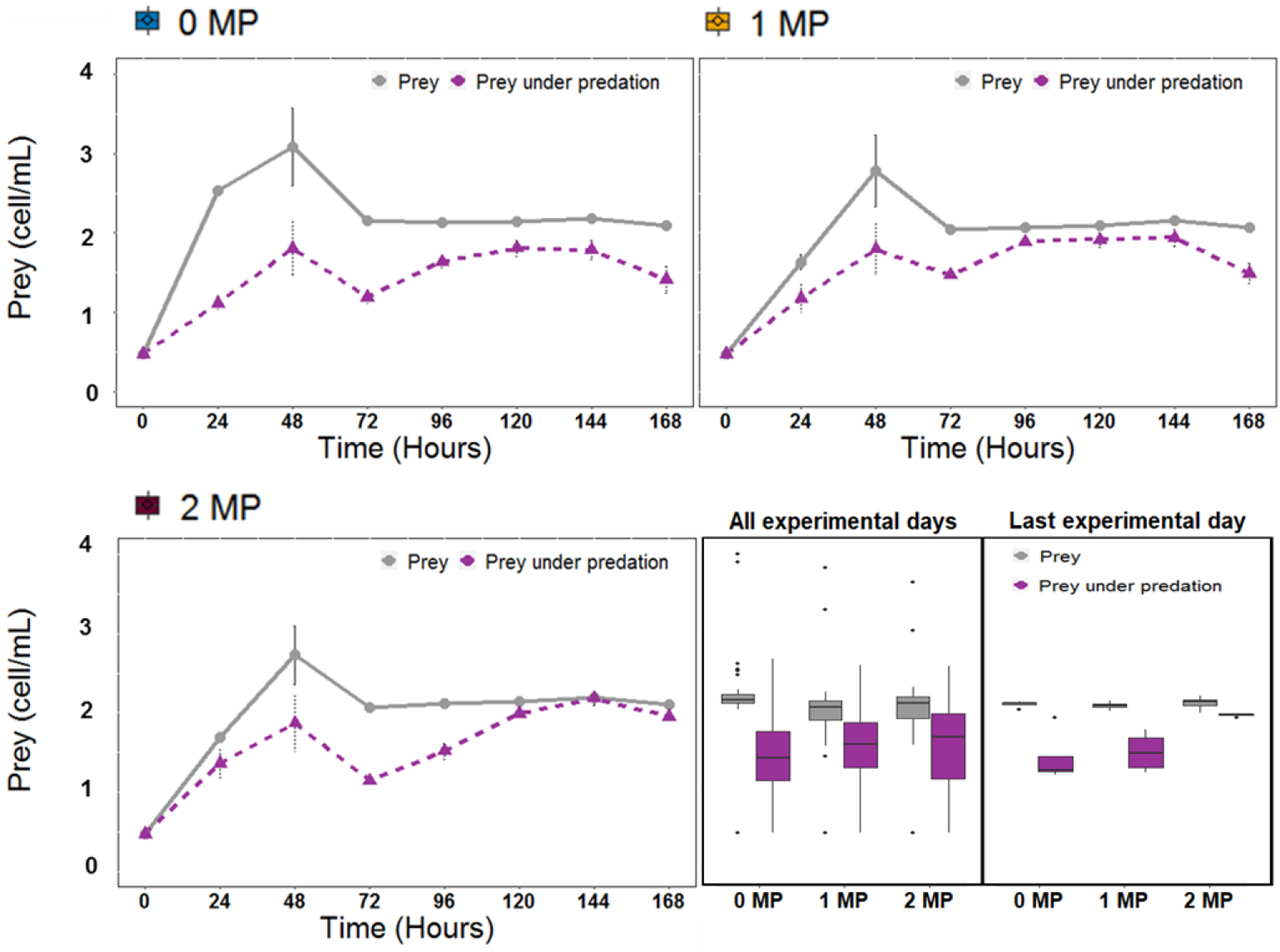
Prey population size (10^9^) dynamics over time, prey average density of all experimental days by MP treatment and prey density in the last experimental day density by MP treatment in single population and in communities in the absence and presence of MP-Fe_dis_ released from the three magnetic particles concentration tested (0 MP, 1 MP and 2 MP).

To further compare the effects of MP-Fe_dis_ on the community level, we tested for differences in predator-prey ratios as an estimate for species interaction strength (Fig. 3); higher ratios indicate stronger predation pressure on the prey. MP treatments were a source of variation between the communities resulting in temporal peaks of higher predator-prey rations in communities treated with the higher MP treatment but resulted on similar communities over time (GEE, MP: W=1.69×10^19^, df=2, p<2×10^−16^, MP x Time: W=-4,41×10^18^, df=2, p=1; Time: W=-1.8×10^17^, df=2, p=1; Fig. 3). This result of increasing predation pressure at early stages of MP exposure, is in line with the observations in the single species during a period of 48 hours. In single species experiments, we found a trend towards higher predator densities in the MP treatment compared to 0 MP treatments (Fig. 1a). However, over time we observed similar ratios for the different MP treatments, and thus similar community responses after few days from the exposure event. Nonetheless, differences in predation pressure among MP treatments resulted in a delay in the higher predation pressure peak in 1 MP and 2 MP treatment compared to the 0 MP treatment. This temporal variation in predation pressures might influence species interactions driving diverse population dynamics and evolutionary responses. Specifically, we observed that over time the predator/prey dynamics were similar, however, they might be driven by diverse evolutionary responses of the prey. For instance, in 0 MP the peak of predation pressure occurred after 24 hours of exposure, however in 1 MP and 2 MP treatments it occurred after 48 and 72 hours respectively. The MP exposure might impose a detrimental effect on predator density delaying the peak of predation to a longer time after exposure. In this regard, theory and empirical data show how the perturbation co-occurence, perturbation timing and strength influence coadaptation and select for difference prey defences (Hiltunen et al. 2018; Raatz et al. 2019). Therefore, the observed temporal variation in predation pressures might select for different prey defences in the microbial community.

**Fig. 3.**
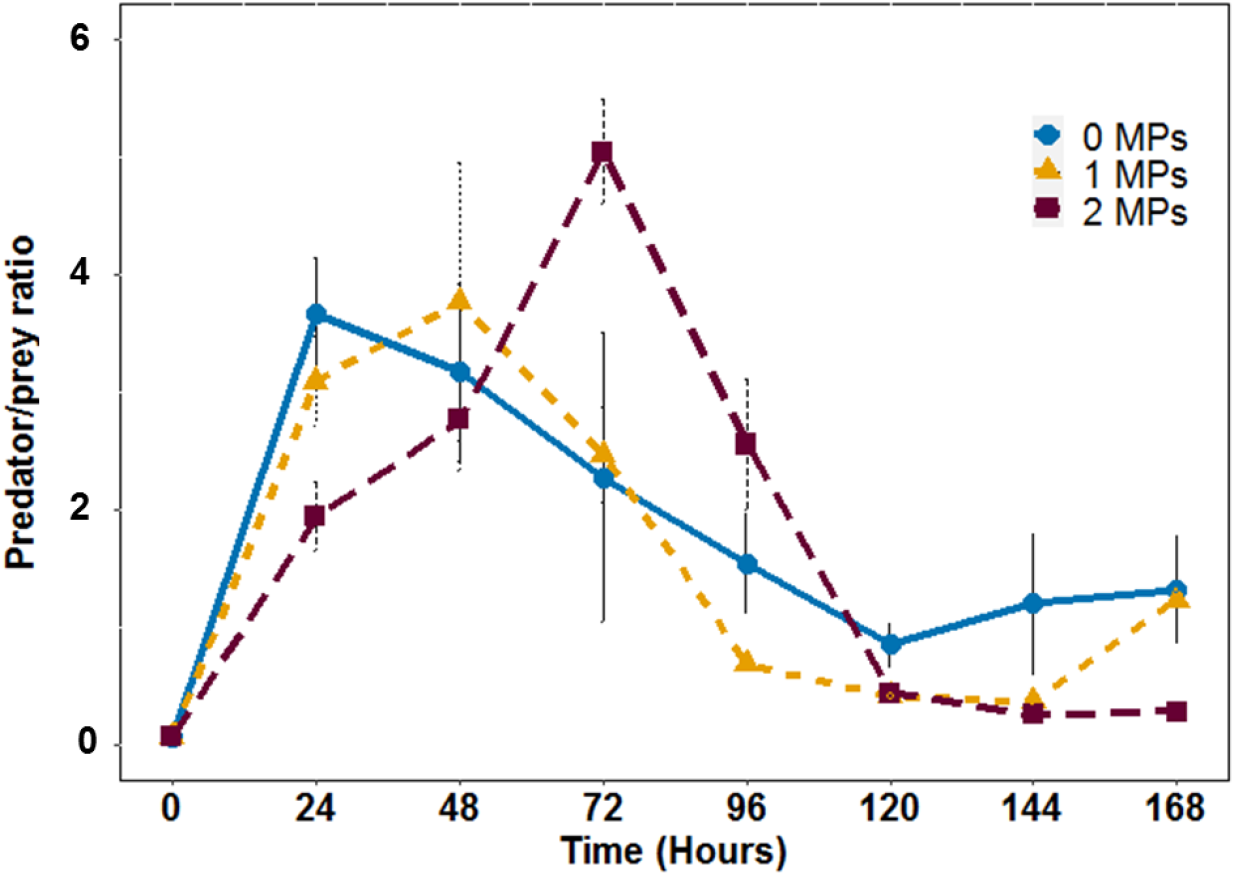
Predator/prey ratios (10^−5^) from the predator-prey community experiment over time at different MP-Fe_dis_ released from the three magnetic particles concentration tested (0 MP, 1 MP and 2 MP). Error bars represent the standard deviations of the mean (mean ± s.d., *n*=4).

As evolution of prey defences can alter predator-prey interactions (Yoshida et al. 2003; Hiltunen et al. 2015) and thus population dynamics, we followed the evolutionary response of the prey to MP treatment (i.e. heritable phenotypic defence trait changes) in the presence and absence of the predator. For this we measured growth rates of the ancestral predator (naïve predator) when grown on same densities of either ancestral prey (naïve prey) or isolated prey (potentially evolved) taken from each transfer of the experiment. From this, we calculated the defence level *D* with zero meaning that the evolved prey had a similar level of defence as the ancestor; values close to 1 indicate a high level of defence compared to the ancestor. As all phenotypic measurements were performed using isolates from the experiment, which grew at least 10 generations in the absence of the MP-Fe_dis_ and the ciliates under standardize conditions before the phenotypes were measured, all observed defence changes are likely genetically determined and not environmentally induced.

Defence evolved but differently in all MP treatments. We found differences for the interaction of time-MP treatment (GEE, MP x Time: W=-4.63×10^27^, df=1, p<2×10^−16^, Fig. 4), for prey defence evolution levels for the different MP treatments (GEE, MP: W=1.07×10^29^, df=2, p<2×10^−16^, Fig. 4) and over time (GEE, Time: W=1.6×10^129^, df=1, p<2×10^−16^, Fig. 4). Specifically, defence evolved to higher level of defence compared to ancestral prey in 0 MP treatment and kept at similar levels over time. However, defence in 1 MP and 2 MP fluctuated over the course of the experiment being: a) always lower compared to defence levels of prey from 0 MP, and b) sometimes similar or even lower than ancestral prey. In addition, other prey traits (e.g. motility) and nutritional quality (e.g. palatability) might influence predation strength, so then, defence levels. In fact, Cairns et al. (2017) studied genomic evolution of the bacterium *Pseudomonas fluorescens* under antibiotics and phage selection pressures alike the metal and predation selection pressure of the here presented microcosmos experiments. They found, genetic evolution related to motility, nutrient transport and metabolic pathways.

**Fig. 4.**
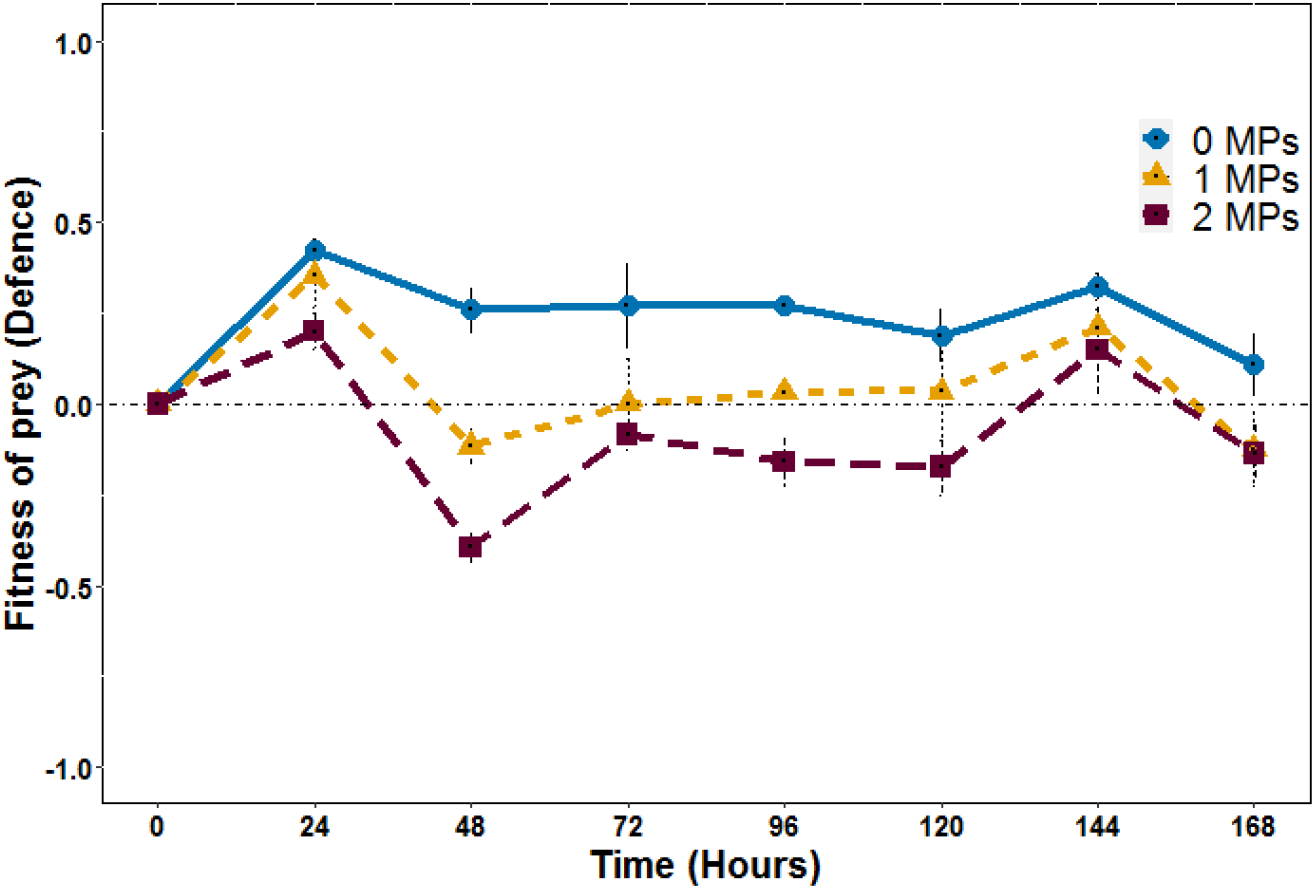
Prey defence dynamics on the predator and prey communities at different MP-Fe_dis_ released from the three magnetic particles concentration tested (0 MP, 1 MP and 2 MP). Dashed horizontal line at the value of 0 is a reference line representing no evolution of defence compared with the ancestors. Error bars represent the standard deviations of the mean (mean ± s.d., *n*=4).

We have assessed the effects of a chemical exposure of MP-Fe_dis_ (coming from MP concentrations intended to be used in lake restoration) on ecological and evolutionary responses in a prey-predator system. In single population, we observed MP-Fe_dis_ favoured an increase of predator_;_ while, prey, after an initial detrimental effect, growth to equal population sizes independently of the MP-Fe_dis_ exposure. In the predator-prey community, we observed similar community responses despite of the different MP concentrations. However, the underlying processes that drove the population size changes and ratios over time might differ with and without MP due to prey defence trait evolution. Prey defence evolved differently in the absence *vs*. presence of MP and had a significant effect on the predator’s population growth rate. Prey defence evolution fluctuated in the presence of MP resulting in similar or lower defence levels that ancestral prey suggesting that predator’s population growth was mostly driven by prey abundances in the community. Thus, although we observed similar predator-prey ratios and dynamics over time in all treatments, our results suggest that the relative roles of ecology and evolution differed significantly depending on the presence or absence of MP. In agreement with our findings, other studies on diverse taxa (rotifer-algae, phage-bacteria, protozoan-mosquito, plant-insect) report population dynamics driven by changes where ecology and evolutionary roles vary in response to both abiotic and biotic changes (Yoshida et al. 2007; Becks et al. 2010; TerHorst et al. 2010; Agrawal et al. 2013).

The fact that the interplay of ecological and evolutionary changes are challenging to identify at community levels raises concern for current ecological risk assessments. Understanding how and when evolutionary changes have the potential to feedback onto ecological changes, and *vice versa*, will enrich the interpretation of of risk assessments by identifying the mechanisms driving community changes (e.g. species sorting vs. genetic sorting). We advocate for more experiments considering the relevance of ecological and evolutionary changes at same time scales under environmentally realistic scenarios. It will provide crucial information for understanding how human-driven selection shapes communities and improve ecological risk assessments.

## Author’s contributions

AdA conceived the study, AdA, LB, IdV design of the study; AdA carried out the experiment, AdA, LB analysed the data; AdA, LB wrote the manuscript, all authors edited the manuscript.

## Acknowledgments

We thank Goekce Ayan for sharing valuable protocols and experience with the experimental system. This work was supported by Junta de Andalucía projects P10-RNM-6630 and P11-FQM-7074 (Proyectos de Excelencia, Spain), MINECO CTM 2013-46951-R, MAT 2013-44429-R and PCIN 2015-051 projects (Spain) and by the European Regional Development Fund (ERDF).

## References

Agrawal AA, Johnson MTJ, Hastings AP, Maron JL (2013) A Field Experiment Demonstrating Plant Life-History Evolution and Its Eco-Evolutionary Feedback to Seed Predator Populations. The American Naturalist 181:S35–S45. https://doi.org/10.1086/666727

Álvarez-Manzaneda I, Guerrero F, del Arco AI, et al (2019) Do magnetic phosphorus adsorbents used for lake restoration impact on zooplankton community? Science of The Total Environment 656:598–607. https://doi.org/10.1016/j.scitotenv.2018.11.375

Ayan G (2018) Using experimental evolution to evaluate diversification of Pseudomonas fluorescens SBW25 in complex environments. https://macau.uni-kiel.de/receive/diss_mods_00022674

Becks L, Ellner SP, Jones LE, Hairston Nelson G.JG (2010) Reduction of adaptive genetic diversity radically alters eco-evolutionary community dynamics. Ecology Letters 13:989–997. https://doi.org/10.1111/j.1461-0248.2010.01490.x

Beketov MA, Liess M (2006) The influence of predation on the chronic response of Artemia sp. populations to a toxicant. Journal of Applied Ecology 43:1069–1074. https://doi.org/10.1111/j.1365-2664.2006.01226.x

Bell G (2017) Evolutionary Rescue. Annu Rev Ecol Evol Syst 48:605–627. https://doi.org/10.1146/annurev-ecolsys-110316

Bell G, Gonzalez A (2011) Adaptation and evolutionary rescue in metapopulations experiencing environmental deterioration. Science 332:1327–1330. https://doi.org/10.1126/science.1203105

Brady SP, Monosson E, Matson CW, Bickham JW (2017) Evolutionary toxicology: Toward a unified understanding of life’s response to toxic chemicals. Evolutionary Applications 10:745–751. https://doi.org/10.1111/eva.12519

Brooks AC, Gaskell PN, Maltby LL (2009) Sublethal effects and predator-prey interactions: implications for ecological risk assessment. Environmental toxicology and chemistry / SETAC 28:2449–2457. https://doi.org/10.1897/09-108.1

Cairns J, Frickel J, Jalasvuori M, et al (2017) Genomic evolution of bacterial populations under coselection by antibiotics and phage. Molecular Ecology 26:1848–1859. https://doi.org/10.1111/mec.13950

Carpenter SR (2008) Phosphorus control is critical to mitigating eutrophication. PNAS 105:11039– 11040

Chiadò A, Varani L, Bosco F, Marmo L (2013) Opening Study on the Development of a New Biosensor for Metal Toxicity Based on Pseudomonas fluorescens Pyoverdine. Biosensors 3:385–399. https://doi.org/10.3390/bios3040385

Dayeh VR, Lynn DH, Bols NC (2005) Cytotoxicity of metals common in mining effluent to rainbow trout cell lines and to the ciliated protozoan, Tetrahymena thermophila. 19:399–410. https://doi.org/10.1016/j.tiv.2004.12.001

De Meester L, Brans KI, Govaert L, et al (2019) Analysing eco-evolutionary dynamics—The challenging complexity of the real world. Functional Ecology 33:43–59. https://doi.org/10.1111/1365-2435.13261

de Vicente I, Merino-Martos A, Cruz-Pizarro L, de Vicente J (2010) On the use of magnetic nano and microparticles for lake restoration. Journal of Hazardous Materials 181:375–381. https://doi.org/10.1016/j.jhazmat.2010.05.020

del Arco A, Álvarez-Manzaneda I, Funes A, et al (2021) Assessing the toxic effects of magnetic particles used for lake restoration on phytoplankton: A community-based approach. Ecotoxicology and Environmental Safety 207:111288. https://doi.org/10.1016/j.ecoenv.2020.111288

del Arco A, Parra G, de Vicente I (2017) Going deeper into phosphorus adsorbents for lake restoration: Combined effects of magnetic particles, intraspecific competition and habitat heterogeneity pressure on Daphnia magna. Ecotoxicology and environmental safety 148:513–519. https://doi.org/10.1016/j.jmr.2004.05.021

del Arco AI, Parra G, Rico A, Van den Brink PJ (2015) Effects of intra- and interspecific competition on the sensitivity of aquatic macroinvertebrates to carbendazim. Ecotoxicology and Environmental Safety 120:27–34. https://doi.org/10.1016/j.ecoenv.2015.05.001

Development Core Team R A Language and Environment for Statistical Computing. R Foundation for Statistical Computing. http://www.r-project.org/

Friman VP, Jousset A, Buckling A (2014) Rapid prey evolution can alter the structure of predator-prey communities. Journal of Evolutionary Biology 27:374–380. https://doi.org/10.1111/jeb.12303

Funes A, de Vicente J, Cruz-Pizarro L, et al (2016) Magnetic microparticles as a new tool for lake restoration: A microcosm experiment for evaluating the impact on phosphorus fluxes and sedimentary phosphorus pools. Water Research 89:366–374. https://doi.org/10.1016/j.watres.2015.11.067

Gibbs MM (1979) A simple method for the rapid determination of iron in natural waters. Water Research 13:295–297. https://doi.org/10.1016/0043-1354(79)90209-4

Gibson B, Wilson DJ, Feil E, Eyre-Walker A (2018) The distribution of bacterial doubling times in the wild. Proceedings of the Royal Society B: Biological Sciences 285:. https://doi.org/10.1098/rspb.2018.0789

Halekoh U, Højsgaard S, Yan J (2006) The R Package geepack for generalized estimating equations. Journal of Statistical Software 15:1–11. https://doi.org/10.18637/jss.v015.i02

Hendry AP, Kinnison MT, Heino M, et al (2011) Evolutionary principles and their practical application. 4:159–183. https://doi.org/10.1111/j.1752-4571.2010.00165.x

Hiltunen T, Ayan GB, Becks L (2015) Environmental fluctuations restrict eco-evolutionary dynamics in predator-prey system. Proc R Soc B 2 282:1–7. https://doi.org/10.1098/rspb.2015.0013

Hiltunen T, Becks L (2014) Consumer co-evolution as an important component of the eco-evolutionary feedback. Nature Communications 6:1–8. https://doi.org/10.1038/ncomms6226

Hiltunen T, Cairns J, Frickel J, et al (2018) Dual-stressor selection alters eco-evolutionary dynamics in experimental communities. Nat Ecol Evol 2:1974–1981. https://doi.org/10.1038/s41559-018-0701-5

Jeppensen E, Kristensen P, Jensen JP, et al (1991) Recovery resilience following a reduction in external phosphorus loading of shallow, eutrophic Danish lakes: duration, regulating factors and methods for overcoming resilience. Ecosystem research in freshwater environment recovery 48:127–148

Kassen R (2014) Experimental Evolution and the Nature of Biodiversity

Koch U, von Elert E, Straile D (2009) Food quality triggers the reproductive mode in the cyclical parthenogen Daphnia (Cladocera). Oecologia 159:317–324. https://doi.org/10.1007/s00442-008-1216-6

Lindsey HA, Gallie J, Taylor S, Kerr B (2013) Evolutionary rescue from extinction is contingent on a lower rate of environmental change. Nature 494:463–467. https://doi.org/10.1038/nature11879

Lopez Pascua L, Gandon S, Buckling A (2012) Abiotic heterogeneity drives parasite local adaptation in coevolving bacteria and phages. Journal of Evolutionary Biology 25:187–195. https://doi.org/10.1111/j.1420-9101.2011.02416.x

Madan NJ, Marshall WA, Laybourn-Parry J (2005) Virus and microbial loop dynamics over an annual cycle in three contrasting Antarctic lakes. Freshwater Biology 50:1291–1300. https://doi.org/10.1111/j.1365-2427.2005.01399.x

Madoni P, Romeo MG (2006) Acute toxicity of heavy metals towards freshwater ciliated protists. 141:1–7. https://doi.org/10.1016/j.envpol.2005.08.025

Matthews B, Narwani A, Hausch S, et al (2011) Toward an integration of evolutionary biology and ecosystem science. Ecology Letters 14:690–701. https://doi.org/10.1111/j.1461-0248.2011.01627.x

Matz C, Kjelleberg S (2005) Off the hook - How bacteria survive protozoan grazing. Trends in Microbiology 13:302–307. https://doi.org/10.1016/j.tim.2005.05.009

Merino-Martos A, de Vicente J, Cruz-Pizarro L, de Vicente I (2011) Setting up High Gradient Magnetic Separation for combating eutrophication of inland waters. Journal of Hazardous Materials 186:2068–2074. https://doi.org/10.1016/j.jhazmat.2010.12.118

Murdoch WW, Briggs CJ, Nisbet RM (2003) Consumer-Resource Dynamics. Princeton University Press, New Jersey, USA

Palkovacs EP, Hendry AP (2010) Eco-evolutionary dynamics: intertwining ecological and evolutionary processes in contemporary time. F1000 biology reports 2:. https://doi.org/10.3410/B2-1

Pohlert T (2014) The Pairwise Multiple Comparison of Mean Ranks Package (PMCMR). R package. http://CRAN.R-project.org/package=PMCMR.

Post DM, Palkovacs EP (2009) Eco-evolutionary feedbacks in community and ecosystem ecology: interactions between the ecological theatre and the evolutionary play. Phil Trans R Soc B 364:1629–1640. https://doi.org/10.1098/rstb.2009.0012

Raatz M, Velzen E, Gaedke U (2019) Co‐adaptation impacts the robustness of predator–prey dynamics against perturbations. Ecol Evol 9:3823–3836. https://doi.org/10.1002/ece3.5006

Ramsayer J, Kaltz O, Hochberg ME (2013) Evolutionary rescue in populations of Pseudomonas fluorescens across an antibiotic gradient. Evolutionary Applications 6:608–616. https://doi.org/10.1111/eva.12046

Sentis A, Gémard C, Jaugeon B, Boukal DS (2017) Predator diversity and environmental change modify the strengths of trophic and nontrophic interactions. Global Change Biology 23:2629–2640. https://doi.org/10.1111/gcb.13560

Smith VH, Schindler DW (2009) Eutrophication science: where do we go from here? Trends in Ecology and Evolution 24:201–207. https://doi.org/10.1016/j.tree.2008.11.009

Søndergaard M, Kristensen P, Jeppesen E (1993) Eight years of internal phosphorus loading and changes in the sediment phosphorus profile of Lake Søbygaard, Denmark. 253:345–356

Straub L, Strobl V, Neumann P (2020) The need for an evolutionary approach to ecotoxicology. Nature Ecology & Evolution 4:895–895. https://doi.org/10.1038/s41559-020-1194-6

TerHorst CP, Miller TE, Levitan DR (2010) Evolution ofprey in ecological time reduces the effect size ofpredators in experimental mesocosms. Ecology 91:629–636

Van den Brink PJ, Klein SL, Rico A (2017) Interaction between stress induced by competition, predation, and an insecticide on the response of aquatic invertebrates. Environmental Toxicology and Chemistry 36:2485–2492. https://doi.org/10.1002/etc.3788

Wang F, Yao J, Chen H, et al (2014) Evaluate the heavy metal toxicity to Pseudomonas fluorescens in a low levels of metal-chelates minimal medium. Environmental science and pollution research international 21:9278–86. https://doi.org/10.1007/s11356-014-2884-x

Worden AZ, Not F (2008) Ecology and Diversity of Microorganisms. In: Kirchman DL (ed) Microbial Ecology of the Oceans, Second Edi. John Wiley & Sons, Inc, pp 159–206

Workentine ML, Harrison JJ, Stenroos PU, et al (2008) Pseudomonas fluorescens’ view of the periodic table. Environmental Biology 10:238–250. https://doi.org/10.1111/j.1462-2920.2007.01448.x

Yoshida T, Ellner SP, Jones LE, et al (2007) Cryptic population dynamics: Rapid evolution masks trophic interactions. PLoS Biology 5:1868–1879. https://doi.org/10.1371/journal.pbio.0050235

Yoshida T, Jones LE, Ellner SP, et al (2003) Rapid evolution drives ecological dynamics in a predator – prey system. Letters to nature 424:303–306

